# Influence of culturing media components on the growth and microbial induced calcium carbonate precipitation (MICP) activity of *Lysinibacillus sphaericus*

**DOI:** 10.1101/2022.05.23.493178

**Authors:** Seyed Ali Rahmaninezhad, Yaghoob A. Farnam, Caroline L. Schauer, Ahmad Raeisi Najafi, Christopher M. Sales

## Abstract

In order to identify appropriate environmental conditions and media components that are either essential or that enhance its growth and Microbial induced calcium carbonate precipitation (MICP) activity, in this study, a series of experiments were conducted to investigate the effects of media components and oxygen conditions on the growth rate and MICP activity of *Lysinibacillus sphaericus* strain MB 284. From these experiments, it was observed that aerobic conditions could lead to increased calcium carbonate production and up to three times faster growth rates by strain MB284 when compared to anoxic conditions. It was also determined that considering the measured growth rate, final biomass concentration, ureolysis activity, amount of calcium carbonate precipitation, and cost of media components for designing undefined culture media for industrial applications, yeast extract is the most economically appropriate option. In our attempts to grow strain MB284 in urea, sucrose, and ammonium acetate as its sole carbon source in minimal media, it was observed it is auxotrophic and that casamino acids and casein are essential for its growth. Even though our experiments agree with the literature that the addition of urea enhances the growth and MICP activity of *L. sphaericus*, it was discovered that when the initial urea concentration was greater than 3 g/l, the growth rate of strain MB284 can be temporarily inhibited until enough cells and urease are produced. These results reveal that the growth and MICP activity of strain MB284 during its application for bio-self healing can be highly dependent on environmental and nutrient conditions.

**Importance:**

- Aerobic conditions increase calcium carbonate production by strain MB284
- yeast extract is the most economically appropriate option for industrial applications of MICP
- *Lysinibacillus sphaericus* strain MB 284 is auxotroph and casamino acids and casein are essential for its growth
- the growth rate of strain MB284 can be temporarily inhibited with urea

## 1. Introduction

Pathways for microbial induced calcium carbonate precipitation (MICP) process differ primarily based on the type of electron donors and acceptors used and include, but are not limited to, urea hydrolysis (ureolysis) [1, 2, 3], ammonification [4], aerobic oxidation [5, 6], denitrification (nitrate reduction) [2], photosynthesis [7], anaerobic sulfide oxidation [8], and methane oxidation [9]. A number of cultures and organisms can perform MICP, with varying degrees of calcium carbonate mineralization efficiency, and include autotrophs such as *Cyanobacteria, Synechococcus*, and *Prochlorococcus* and heterotrophs such as *Sporosarcina pasteurii* (*Bacillus sphaericus*), *B. megaterium, B. subtilis, B*.*cereus, B. cohnii, B. pseudofirmus, B. alkalinitrilicus, Diaphorobacter nitroreducens, Pseudomonas aeruginosa, Desulfovibrio brasiliensis, Desulfovibrio vilgaris, B. mucilaginous* [10, 6]. Among this wide spectrum of axenic and non-axenic cultures that can produce carbonate ions and precipitate calcium carbonate in alkaline conditions, *Lysinibacillus sphaericus*, formerly known as *B. sphaericus*, has abilities that make this species a favorable biological component in MICP activities such as: forming long-lasting spores even more than 50 years in environmentally harsh conditions [6], creating rhombohedral and tightly crystal-packed layers, having high rates of urea hydrolysis and calcium carbonate precipitation, and precipitating bio-minerals based on both urea hydrolysis and denitrification pathways [2, 11, 12, 13, 14, 5].

Anbu et al. reported that when *L. sphaericus* 21776 was applied as a bio-agent under aerobic conditions, the kinetics of urea hydrolysis (k_urea_) was 6.2e-8 h^-1^ ml CFU^-1^, while that of was 3.2e-8 h^-1^ ml CFU^-1^ for *Sporosarcina pasteurii* ATCC 11859 [3]. Regarding the urea decomposition percent, different *L. sphaericus* including LMG 22257 [2, 15], LMG 21776 [3], and LMG 22257 [15] could degrade 100% of urea after around one day, while after a similar period, *Sporosarcina psychrophile* DSM 6497 [11], *S. pasteurii* DSM 33 [11] and *S. pasteurii* ATCC 11859 [3] as commonly used species in urea hydrolysis could degrade only 53%, 72%, and 43% of urea, respectively. In addition, *L. sphaericus* possesses high calcium carbonate precipitation kinetics (k_precip_) in comparison with other isolates. For example, the k_precip_ of *L. sphaericus* 21776 was 0.604 h^-1^ [3], while that of for *S. pasteurii* ATCC 11859 was 0.253 h^-1^ [3], 0.014 h^-1^ [16], 0.112 [17], and 0.065 [18] based on different studies and for *S. pasteurii* ATCC 6453 was 0.116 h^-1^ [19] in nearly similar environmental conditions.

Since the availability of oxygen could be limiting under some conditions in concrete [5, 15], it is important the effects of aerobic and anoxic conditions on the *L. sphaericus* growth rate, urea hydrolysis, and calcium carbonate precipitation be investigated. Zhu et al. has already reported that *L. sphaericus* LMG 22257 can precipitate bio-minerals via both urea hydrolysis and denitrification pathways under anoxic conditions [2]. It has been previously reported that the concentration of oxygen has only a marginal effect on urease activity [2, 15] and *L. sphaericus* YS11 is not able to precipitate calcium carbonate under hypoxia (i.e., anaerobic conditions) [13]. Since both biomass concentration and urea hydrolysis are affected by oxygen concentrations and the number of cells (which correlates with biomass concentration) can affect the number of nucleation sites for scavenging divalent cations and subsequently calcium carbonate precipitation, it is important that all of these interconnected parameters be simultaneously evaluated to find the exact role of oxygen in the MICP process.

In addition to examining the effects of oxygen conditions, this study compares the effect of different culture media components on MICP activity and the growth of *L. sphaericus* strain MB284. Although a variety of undefined culture media use nutrient-rich sources such as beef extract, peptone [6], and yeast extract [20] to cultivate other types of bioagents for calcium carbonate precipitation, nearly most of the research studies on *L. sphaericus* have been focused on growing this species in yeast extract [13, 2, 11, 21, 22, 23, 15, 14]. Therefore, this study also aimed to investigate the advantage of growing strain MB284 in yeast extract over other culture media. Lastly, in order to further study the effects of specific media components, such as urea, on the growth and physiology of *L. sphaericus*, this study also attempted to grow strain MB284 in minimal media. Despite the study of White and Lotay [24] on a limited number of *L. sphaericus* strains revealed that only some of them can grow in minimal media, there is no attempt for growing strain MB284 for MICP purposes.

## 2. Material and Methods

### 2.1. Culture preparation

The strain *Lysinibacillus sphaericus* ATCC 13805TM was revived in a culture medium containing beef extract (10 g/l), peptone (10 g/l), sodium chloride (5 g/l), and distilled water. The pH of the culture media was adjusted to 7 with the use of Phosphate Buffered Saline (PBS) and autoclaved at 121 °C for 15 minutes before inoculation. Then samples were stored in the incubator shaker at 35 °C and 125 rpm for around one day. When the optical density of the culture at the wavelength of 600 nm (OD_600_) reached around 1 A, an appropriate amount of culture media containing bacteria was taken and transferred into the appropriate tubes for centrifuging. Pellets were extracted from culture media at 20 °C at 7830 rpm for 10 minutes then were washed three times with PBS solution before they were added into the desired media. We observed the concentration of strain MB284 was around 10^9^ cells/ml when the OD_600_ was 1 A in the culture media that was similar to the *L. sphaericus* LMG 22257 [15].

The analytical grade culture media including peptone, beef, yeast, meat extract, skim milk powder, casamino acids, casein, thiamine, ammonium acetate, ethylene diamine tetra acetic acid (EDTA), ultra-pure urea, and calcium acetate were used to supply essential components for the growth of strain MB284 and production of calcium carbonate. In addition, two minimal defined media, Minimal Salts Medium (MSM) and EZi medium were used in this experiment for providing nutrients. Table 1 indicates the ingredient of MSM and EZi media.

**Table 1:** The ingredients of the MSM and EZi media.

| MSM medium |  |  |  | EZi medium |  |
| --- | --- | --- | --- | --- | --- |
| Salts | Concentration | Salts | Concentration | Salts | Concentration |
| NaNO<sub>3</sub> | 1 g/l | ZnSO<sub>4</sub>*7H<sub>2</sub>O | 5.74 mg/l | KH<sub>2</sub>PO<sub>4</sub> | 0.5 g/l |
| K<sub>2</sub>SO<sub>4</sub> | 0.17 g/l | MnSO<sub>4</sub>* H<sub>2</sub>O | 3.38 mg/l | Na<sub>2</sub>SO<sub>4</sub> | 2 g/l |
| MgSO<sub>4</sub>*7H<sub>2</sub>O | 0.037 g/l | H<sub>3</sub>BO<sub>3</sub> | 1.24 mg/l | KNO<sub>3</sub> | 2 g/l |
| CaSO<sub>4</sub>*2H<sub>2</sub>O | 0.0086 g/l | CoMoO<sub>4</sub>*H<sub>2</sub>O | 0.94 mg/l | MgSO<sub>4</sub>*7H<sub>2</sub>O | 1 g/l |
| KH<sub>2</sub>PO<sub>4</sub> | 0.53 g/l | FeSO<sub>4</sub>*7H<sub>2</sub>O | 0.224 g/l | FeSO<sub>4</sub> | 0.0004 g/l |
| K<sub>2</sub>HPO<sub>4</sub> | 1.06 g/l | H<sub>2</sub>SO<sub>4</sub> | 0.1 ml/l | DI water | 1 liter |
| KI | 1.66 mg/l | CuSO<sub>4</sub>*5H<sub>2</sub>O | 5 mg/l |  |  |
| DI water | 1 liter |  |  |  |  |

### 2.2. Chemical and Biological Analysis Methods

The concentration of ammonia was measured by spectrophotometer (DR 2500, Hack, US). The pH and electric conductivity (EC) of the culture media was measured by pH meter (Pro DSS, US) and EC meter (Mettler Tolido) [25, 26]. For measuring the amount of calcium carbonate precipitation, 30 ml of the solution were transferred into the tube and well mixed for one minute by a mixer. Then, 20 g/l of calcium acetate was added into the tube and again it was stirred by the mixer for one minute. After that, it was stored in an incubator at 30 °C for 24 hours. Then, the supernatant was removed from the sample by centrifuging at 20 °C at 7830 rpm for 15 minutes. Finally, the sample was placed into the desiccator (VWR, incubating orbital shaker) at 50 °C for 1 day. The difference between the initial and final weights indicated the amount of calcium carbonate production. For normalizing the production of calcium carbonate, the obtained value was divided by the maximum amount of OD_600_ of culture media before adding calcium acetate.

The aerobic condition was provided for bacteria by covering the neck of bottles with fitted foam that allows air to easily penetrate the bottles through the foam. However, sample bottles were thoroughly filled with culture media and were completely capped with plastic lids while there was not any headspace between the media and lids to provide anoxic condition. Since there was no headspace in the sample bottles, nitrogen flushing was not done. In addition, Wang et al showed that the *L. sphaericus* strains are able to consume all of the dissolved oxygen shortly after inoculation [15] the culture media did not purge with nitrogen for providing anoxic conditions. All samples were stored in the shaker at 30 °C during the experiment and run in triplicate.

## 3. Results and Discussions

### 3.1 Effect of nutrient-rich sources in undefined media on bacterial growth and MICP activity

A wide variety of undefined nutrient-rich sources has been used for biomineralization, including yeast extract, lactose mother liquor, corn steep liquor, meat extract, milk extract, glucose, Luria broth, tryptic soy broth, carbohydrate, amino acids, sodium acetate, lactate sugarcane molasses, peptone, beef extract, lentil seed power, glucose, calcium lactate, sugar, and beef extract [10, 1, 11, 6, 27, 28, 15]. In this study, we investigated the influence of using 20 g/l of peptone, meat, yeast extract, or skim milk powder as the nutrient-rich source in undefined media on the growth and MICP activity of strain MB284, by comparing the exponential growth rate (µ), maximum and final biomass concentration, the stationary phase length, the EC increment (ΔEC), and amount of calcium carbonate produced across these nutrient-rich sources.

The results from these investigations reveal that no significant difference in the amount of the lag phase and the bacterial growth rate (µ) had been observed in the examined media except skimmed milk powder that had a considerably lower growth rate. The lag phase was less than 10 hours and µ were 1.8, 1.7, 1.7, and 1.6 (OD_600_/day) for beef, yeast, and meat extract, and peptone; respectively. A similar lag phase around 10 hours and a growth rate of around 1.2 (OD_600_/day) have been reported by Zue et al and Wang et al when *L. sphaericus* LMG 22257 had been cultured in yeast extract containing urea [2, 15]. It was also observed in our experiments that the maximum OD_600_ of the cultures grown in yeast extract, meat extract, peptone, and beef extract were relatively equivalent, being 1.73, 1.7, 1.61, and 1.54 A, respectively, but roughly 3 times more than skimmed milk powder. Biomass concentration that is related to OD_600_, in addition to its impacts on carbonate ions production that leads to increased pH, affects the number of provided nucleation sites served to initiate calcium carbonate precipitation [28, 10, 1]. Previous research [29, 28] demonstrated that more calcium ions precipitated when bacteria instead of chemicals increased the pH of the microenvironment that indicating the important role of the biomass concentration in biomineralization. Furthermore, we noticed that when skimmed milk powder was used, the death phase starts on the seventh day, while culture grown in the other nutrient-rich sources remained in the stationary phase beyond 40 days.

In addition to its growth kinetics, the pH of MB284 cultures grown in the different nutrient-rich sources was examined. The production of carbonate ions and also hydrolysis of urea that produce ammonia, increase the pH [10, 1]. Therefore, as is expected, the pH increased from 6.5 to around 10 after 40 hours for all nutrient-rich sources except skimmed milk powder that pH reduced from 6.5 to 5.5. Tan and Ramamurthi [30], and Akali and Yetisemiyen [31] reported that the inoculation of some kinds of *Bacillus* species in the milk powder acidifies the environment due to lactic fermentation and lactic acid formation.

Finally, we observed that when calcium acetate was added to the culture media, the normalized produced calcium carbonate respecting to the maximum biomass concentration was 2.31, 1.82, 1.33, 1.10 (g/30 ml solution/OD_600_) for beef extract, peptone, yeast extract, and meat extract; respectively. The overall results delineate that the growth kinetics and MICP activity of strain MB284 in all examined culture media was pretty close to each other except for skimmed milk powder. As a result, the noticeably lower price of the yeast extract in comparison with other media can be decisive factors for selecting a favorable medium for the industrial applications of the MICP process.

### 3.2 Impact of oxygen on bacterial growth and MICP activity

Other studies have reported that the availability of oxygen can have a significant impact on biomineralization [15]. Therefore, in this study, we investigated the growth kinetics and MICP activity of strain MB284 was grown in nutrient-rich sources under aerobic and anoxic conditions. We observed that while the length of the lag phase (8 to 10 hours) was the same under both aerobic and anoxic conditions, a significant difference was shown during the exponential growth phase. Furthermore, the growth rate during the exponential phase under aerobic conditions for beef, yeast, meat extract, and peptone was observed to be 13, 11, 10, and 8 times higher than under anoxic conditions, respectively, implying that the growth kinetics of strain MB284 is strongly related to the presence of oxygen. In addition, it was found that the rate of bacterial growth was not only faster in aerobic conditions but that the final biomass concentration was 3-4 times higher under aerobic than anoxic conditions.

Although strain MB284 grew faster and to greater cell densities under aerobic conditions, the length of the stationary phase (at least until 40 days) was the same under aerobic and anoxic conditions, suggesting that cells of MB284 under anoxic conditions remain metabolically active. In addition to the bacterial growth kinetics, the MICP activity of strain MB284 was compared under aerobic and anoxic conditions. It was observed that the rate of increase in pH under anoxic conditions was slower than the aerobic condition for the first two days, however, the final pH of the solution was approximately the same under both aerobic and anoxic conditions. Since the hydrolysis of urea and production of ammonia and carbonate ions under aerobic urea hydrolysis and changing nitrate to ammonia during the denitrification and anoxic oxidation mechanism are the main reasons for increasing pH in nutrient-rich media, the lower degree of pH increase under anoxic conditions confirms that MB284 is more metabolically active under aerobic conditions.

Similar to the pH, the results showed that the amount of the EC in the nutrient-rich media increased faster under the aerobic condition in comparison with anoxic conditions. In the MICP activity, in the presence of urea, the increase in the ion strength of the solution is a result of producing polar components such as carbonate ions, and ammonia during urea hydrolysis [1]. Therefore, this result showed that although urea can be degraded both under aerobic and anoxic conditions by MB284, the higher rate in increasing EC showed the favorability of oxygen for MB284 in the MICP process. However, in the opposite of the final values of pH that were nearly the same under both conditions, the final value of EC under aerobic condition was around two times higher than anoxic condition. Since the nitrate up-taking rate was faster under anoxic conditions than the aerobic condition, it is concluded that the nitrate removal under the anoxic condition caused the EC of the solution to not increase as much as the aerobic condition.

In regard to the production of calcium carbonate, the results from our study showed that when MB284 was inoculated into the beef, yeast, meat extract, and peptone, the final amount of calcium carbonate under the aerobic condition was three to four times higher than the anoxic condition. One possible reason for this higher calcium carbonate production is the increased biomass concentrations seen under aerobic conditions—where an increased number of cells in solution would provide more nucleation sites that are favorable for scavenging divalent cations such as Ca^+2^ and Mg^2+^ for producing calcium carbonate [28]. There is some disagreement in the literature regarding the effect of oxygen on MICP activity, with some reporting that aerobic conditions lead to lower calcium carbonate precipitation yields than anoxic conditions [32, 5, 27].

Although our results indicate that aerobic conditions are favorable for enhancing MICP activity in strain MB284, its ability to also perform MICP activity under anoxic conditions makes it a versatile organism for biomineralization. For example, oxygen availability in the deeper parts of cracks in concrete or cement could become limited, making it challenging to rely on obligate aerobes to produce calcium carbonate [5, 15]. Nitrate, on the other hand, should be highly available in concrete, since its solubility at 20 °C is 10^5^ times more than oxygen and because concrete typically has high concentrations of nitrate-containing components such as calcium nitrate [33]. Therefore, even when oxygen becomes limited, strain MB284 will still be capable of performing MICP for filling a crack in bio concrete applications.

### 3.3 Auxotrophic growth of *L. sphaericus* in minimal media

Organisms that are unable to synthesize particular organic compounds that may be required for their growth are often considered auxotrophic. Studying the auxotrophy of *L. sphaericus* is important for identifying if *L. sphaericus* requires any special nutrients to grow in concrete or cement for self-healing applications. To test whether *L. sphaericus* strain MB284 is auxotrophic, experiments were conducted with strain MB284 in two different minimal media, MSM and EZi medium, that contained either urea or ammonium acetate as the sole carbon sources. It was observed in these experiments that MB284 could not grow after 16 days of inoculation in the defined minimal media which confirms the auxotrophic property of MB284 (Figure 1 A). This auxotrophic property of *L. sphaericus* was previously reported in the literature [24, 34, 35] that shows only a few species of *L. sphaericus* could grow on minimal media. It is assumed that the lack of the special kind of vitamins, amino acids, or other components such as biotin, thiamin, glutamic acid, glutamate, purine, pyridoxine, serine, asparagine, alanine, phenylalanine, methionine, acid-hydrolyzed casein, ethylene diamine tetra acetic acid (EDTA), adenine, and guanine are required to be added to the minimal nutritional medium to assist for bacterial growth [24]. Therefore, in this experiment different concentrations of thiamine, EDTA, and casamino acids in addition to 20 g/l of urea were added to the MSM medium under the anoxic condition to investigate the bacterial growth. Results showed that strain MB284 could only grow when the MSM medium was supplemented with casamino acids (Figure 1 C) that consequently, the pH of the solution increased (Figure 1 B).

**Figure 1:**
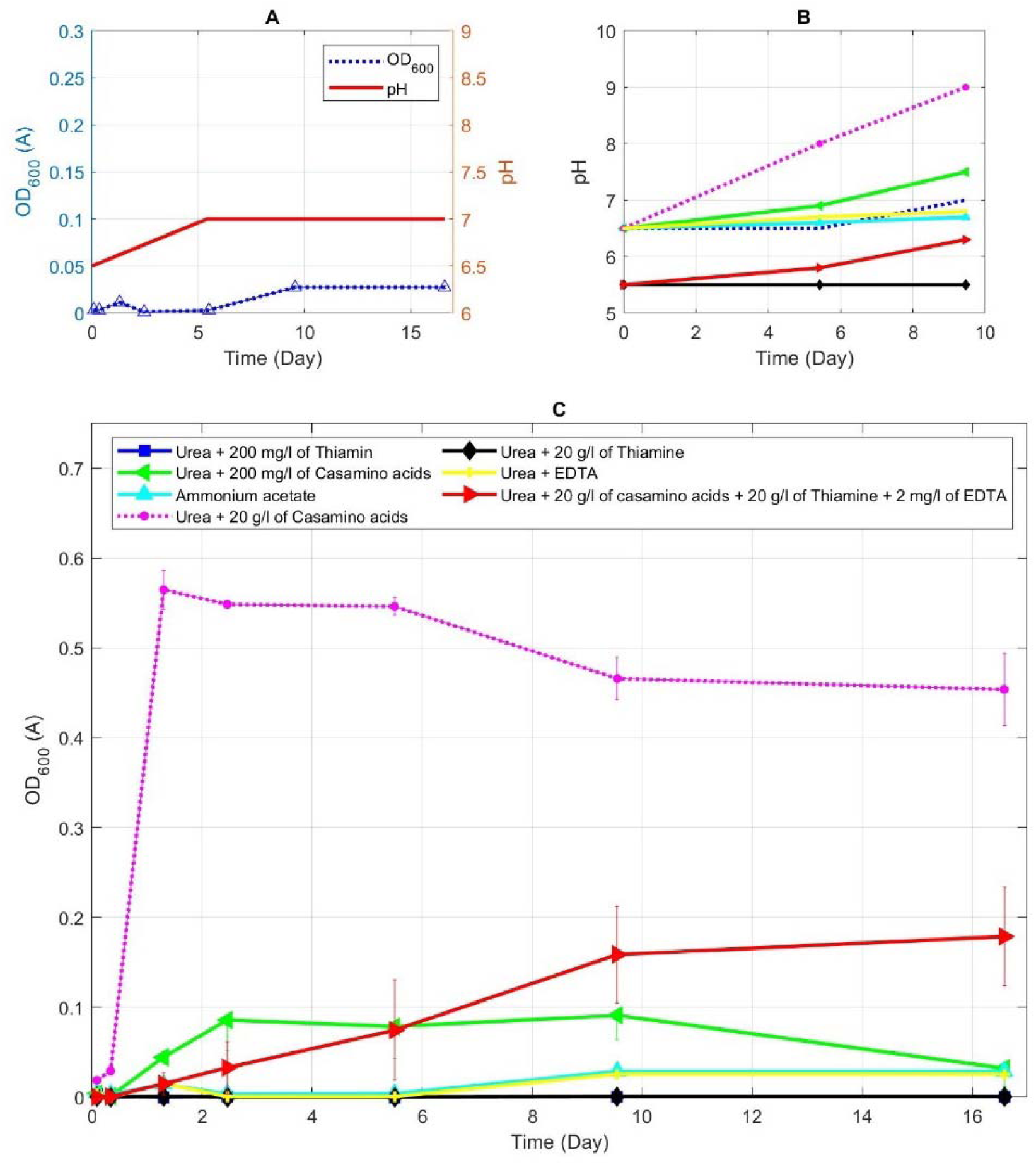
**(A) Investigation of the bacterial growth and pH amount of a culture medium containing MSM and 20 g/l of urea. Investigation of (B) pH changes and (C) the bacterial growth of culture media containing MSM, 20 g/l of urea and different concentrations of thiamin, EDTA, and casamino acids under the anoxic condition**

In order to examine the concentration of casamino acids that was required by strain MB284 for growth, different concentrations of casamino acids were added to the MSM medium containing 20 g/l of urea under the anoxic condition. As Figure 2 A shows, when the initial concentrations of casamino acids are increased, the biomass growth rate and the final biomass concentration increases. However, Figure 2 B illustrates that the slope of the growth rate and final biomass concentration curves significantly decreases when the initial concentration of casamino acids was more than 10 g/l.

**Figure 2:**
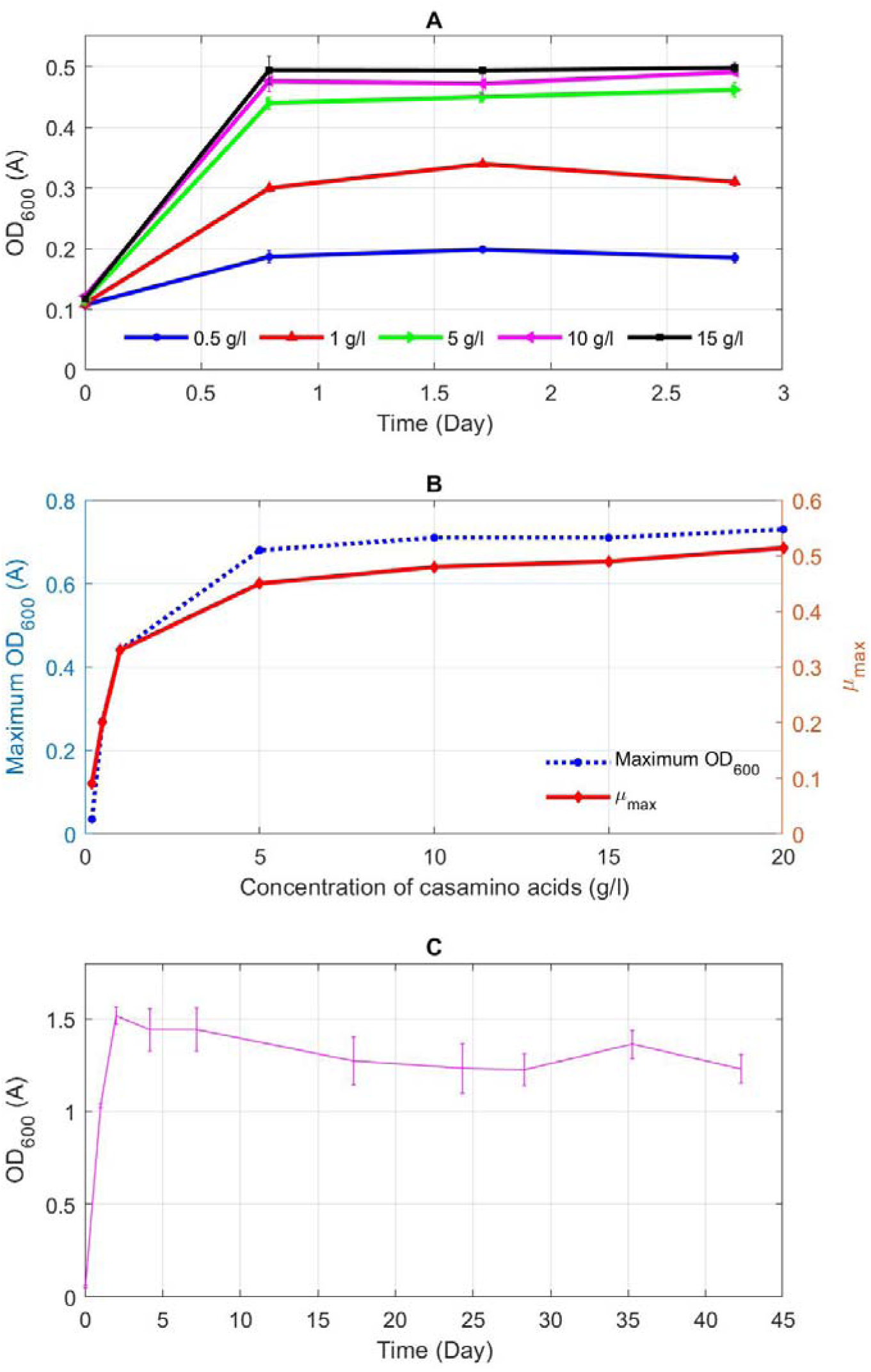
**The effect of adding different concentrations of casamino acids into the MSM containing 20 g/l urea on (A) the bacterial growth curves, (B) maximum biomass concentration and growth rate (µ), in the anoxic condition, and (C) the growth curve of 10 g/l of casamino acid in the aerobic condition**

Further tests were conducted where the concentration of casamino acids was fixed at 10 g/l and 20 g/l of urea was added to the MSM medium to compare growth under the aerobic and anoxic conditions (Figure 2 C). Aligning well with the results of section 3.2, the availability of oxygen triples the biomass concentration in the stationary phase when there is 10 g/l of casamino acids and 20 g/l of urea.

In addition to casamino acids, we also tested the growth of strain MB284 with casein supplemented with the defined minimal media recipes (Figure 3). Casamino acids are derived from casein that have undergone hydrolysis with hydrochloric acid. Casamino acids have been shown to contain highly soluble amounts of alanine, arginine, asparagine, aspartate, cysteine, glutamate, glutamine, glycine, histidine, isoleucine, leucine, lysine, methionine, phenylalanine, proline, serine, threonine, tryptophan, tyrosine, valine, and other peptides [36] that strain MB284 needs for growth in minimal media.

**Figure 3:**
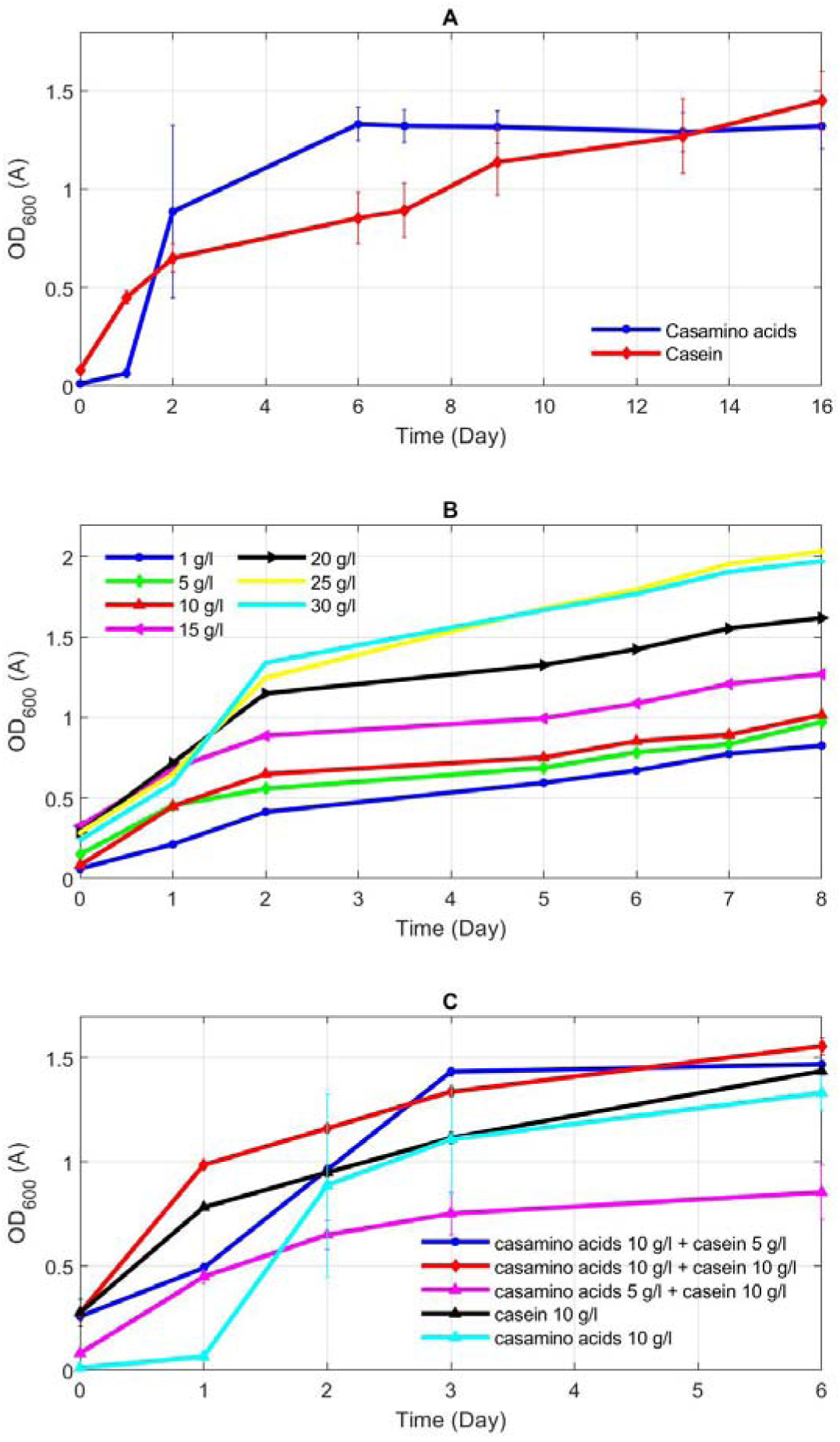
**(A) bacterial growth curve when 10 g/l of casein or casamino acids are added into the MSM medium containing 20 g/l urea; (B) adding different concentration of casein into the MSM medium containing 20 g/l urea; (C) investigating the synchronous effect of adding different mixture of casamino acids and casein concentrations**

As it can be inferred from Figure 3 A, the initial bacterial growth rate in Berekely medium containing casein is considerably higher than that of casamino acids on the first day of the experiment. However, the concentration of biomass became higher in the medium containing casamino acids after the second day and it kept higher values until around day 13^th^. Casein is relatively hydrophobic and poorly soluble in water so, on the opposite of casamino acids, only a small portion of it dissolved right after it was added in minimal medium. Therefore, the initial higher biomass concentration in the medium containing casamino acids can be due to the availability of more required nutrients provided by casamino acids on the first days of the experiment. Based on our observation, although most of the added casein initially into minimal medium remained precipitated, it gradually became completely dissolved as growth was being observed.

Since it was shown that casamino acids and casein had a similar effect on the growth rate of strain MB284 after 16 days, we next investigated the effect of varying the concentration of casein on the growth characteristics of strain MB284. (Figure 3 B). These results show that, while an increase in the initial concentration of casein had a positive effect on enhancing biomass concentration, the growth curves relating to 25 and 30 g/l of casein were very close to each other in the first 8 days of the experiment that indicating the rate of increase in biomass concentration significantly decreased in 30 g/l of casein. Comparing the growth curve of casein and casamino acids shows that in contrast with casamino acids that three distinct growth phases (including lag phase, exponential phase, and stationary phase) can be observed, there was an ascending trend for the growth curve when casein was added into the MSM medium (Figure 3 A). This ascending trend could be a result of the gradually dissolving and accessibility of casein for strain MB284 in the Berekely medium by providing essential growth nutrients.

In addition to evaluating the effects of casamino acids and casein independently on the growth of MB284, we also tested mixtures of different concentrations of both compounds (Figure 3 C). In all combinations of the mixture of casein and casamino acids, lag phases were less than when casamino acids was solely added into the culture medium and the initial growth rate was more than when casein was solely added, suggesting that a combination of casamino acids and casein provides a more favorable culture medium for bacterial growth of MB284.

Lastly, experiments comparing the growth of MB284 in MSM and EZi media and distilled water (DI) containing 10 g/l of casamino acids with or without urea (not shown) illustrated that growth of MB284 only requires casamino acids (or casein) for growth. However, a closer examination of the growth curves of MSM and EZi media with DI water indicates that the addition of minerals has a positive effect on MB284 growth rate in the exponential phase and the amount of the maximum biomass concentration in the first two weeks of the experiment (Figure 4). Since the bacterial growth rate for both MSM and EZi media was nearly identical, it is concluded that the common ions in these two media including sodium, potassium, magnesium, phosphate, and sulfate have more enhancing impacts on the MB284 growth rate. In addition, considering the concentration of salt components there was in the EZi and MSM media showed that the higher concentration of minerals in the MSM medium does not have a significant influence on the MB284 growth curve.

**Figure 4:**
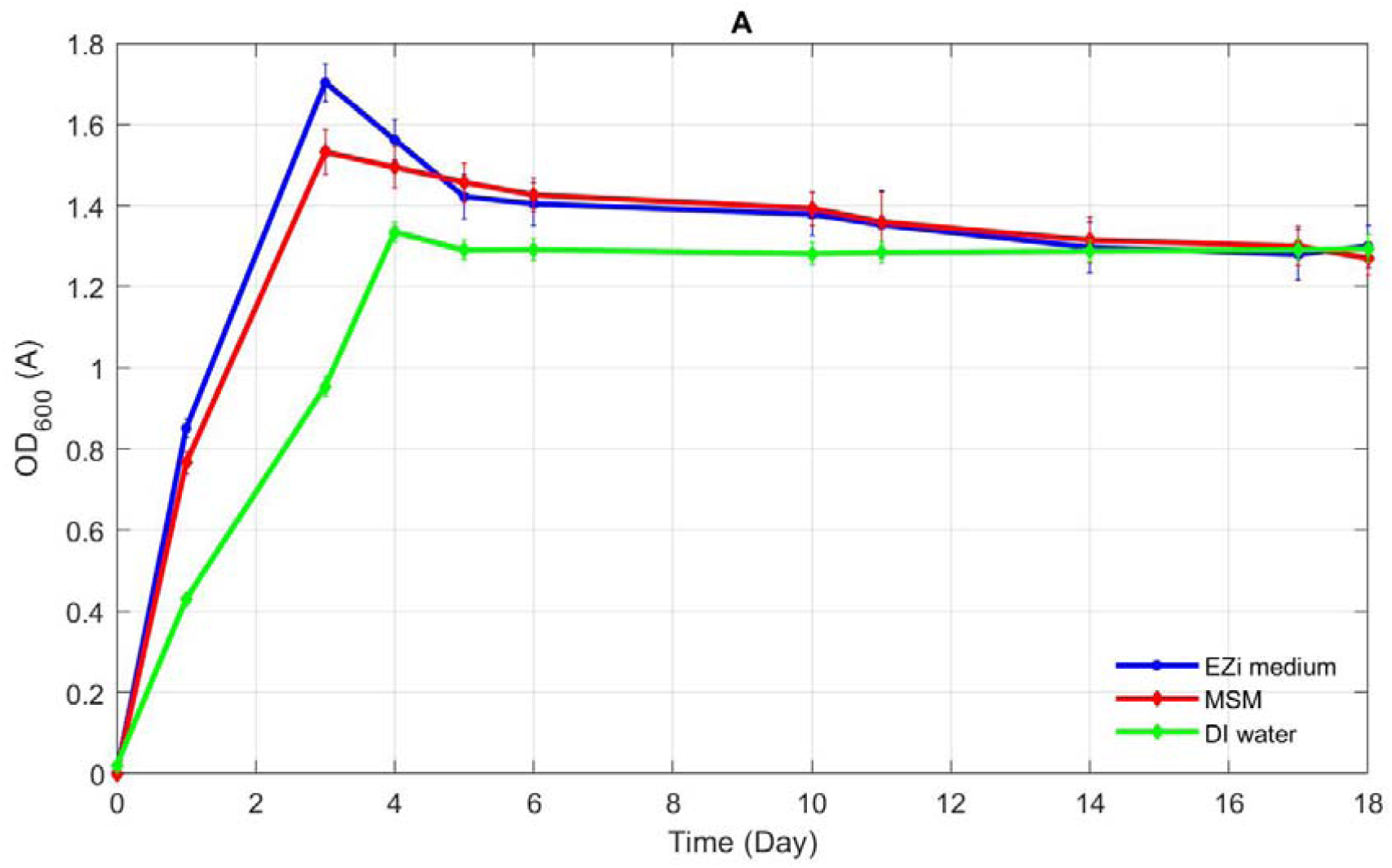
inoculation of *L. sphaericus* into the MSM medium, EZi medium, and DI water in addition to 20 g/l of urea and 10 g/l of casamino acids which are added to all culture media.

### 3.4 The impact of urea on growth and MICP activity

Urea hydrolysis is key for MICP activity because it leads to the conversion of urea to ammonia and CO_2_, which together helps drive calcium carbonate production due to the fact that the increase in ammonia leads to alkaline (i.e., increased pH) conditions that consequentially leads to conversion of CO_2_ to carbonate ions [20, 14, 2, 11]. Biomineralization of calcium carbonate can occur without urea hydrolysis as long as there are sufficient carbonate ions and calcium present and, thus, metabolism of organic compounds by strain MB284 in the absence of urea should lead to calcium carbonate precipitation. Our experimental results showed this to be true, in the absence of urea, strain MB284 was able to produce calcium carbonate when casamino acids, peptone, beef, yeast, or meat extract without the addition of urea. However, we did find that adding 20 g/l of urea to these culture media increased the final biomass concentration (5 to 10%), pH of the solution (30 to 40%), ΔEC (20 to 32%), and the amount of calcium carbonate production (20 to 43%).

Although we observed a positive effect of adding urea on bacterial metabolism and the MICP activity, further experiments revealed that 20 g/l of urea decreased the biomass concentration of strain MB284 in comparison with culture media which was free of urea for the first 5 days of growth (Figure 5 A). These results indicate urea is a growth inhibitor of strain MB284 during the initial growth phase and lead us to investigate the effect of urea concentrations on bacterial growth inhibition (Figure 5 B). The results from this investigation showed that when initial urea concentrations are below 10 g/l the strain MB284 grows faster during the first 3 days of growth following inoculation, which is similar to the results by Bhat et al. [37] that showed the presence of high concentrations of urea can increase the length of the lag phase of growth for mixed microbial consortia originating from loamy soils.

**Figure 5:**
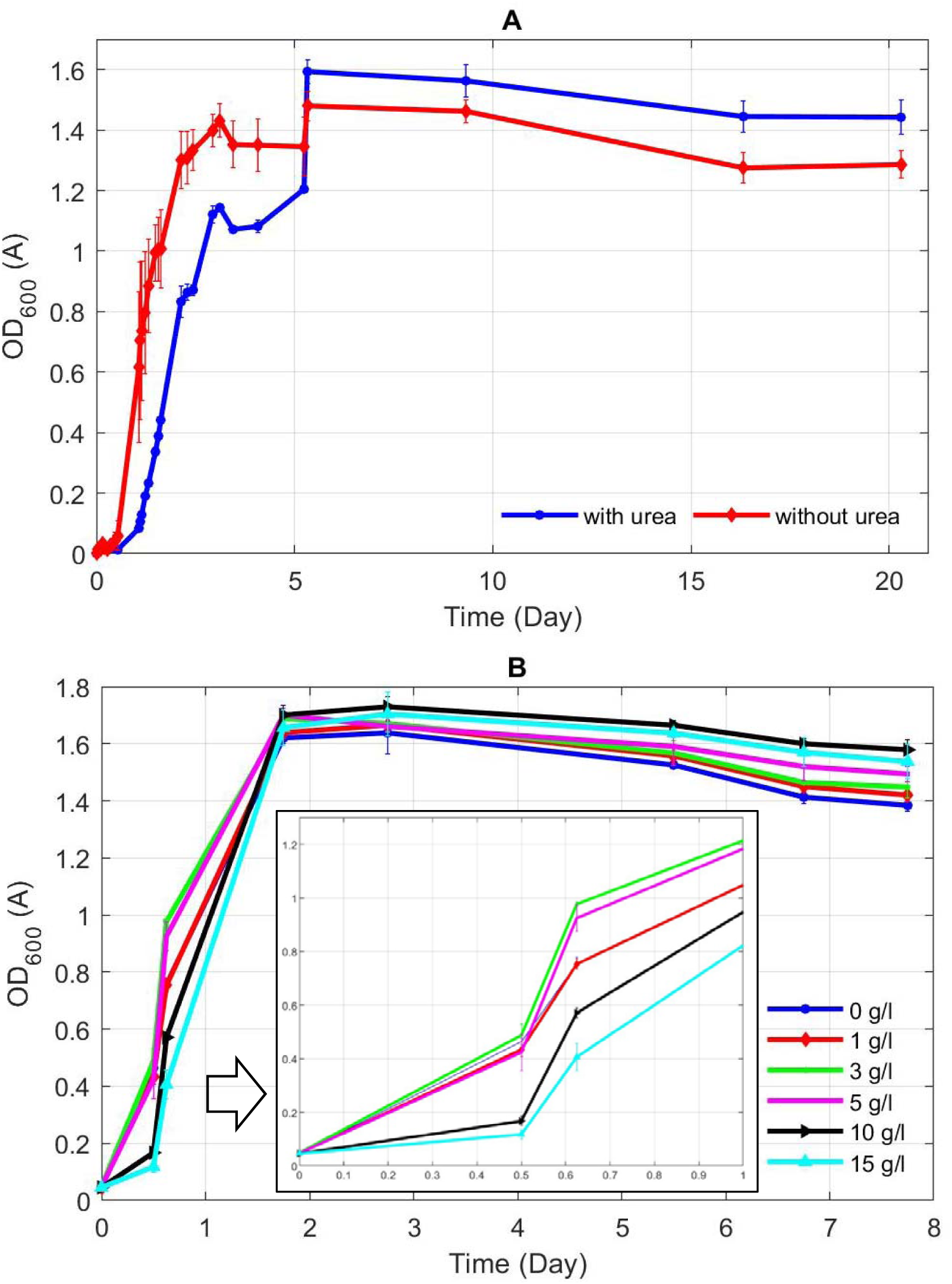
**A: The inhibition effect of 20 g/l of urea on the bacterial growth curve when it is added to MSM medium in addition to 10 g/l of casamino acids, B: The effect of bacterial growth on the different concentrations of urea in the MSM medium and 10 g/l casamino acids**

After this initial 3 day period of growth, it was observed that biomass concentrations of strain MB284 began to increase based on initial urea concentrations, 10 and 15 g/l of urea having the great final biomass concentration. Although Wang et al. [15] reported that urea inhibition occurs with *L. sphaericus* LMG22257 in all tested urea concentrations between 12 to 180 g/l, when lower concentrations of urea was applied, we found that inhibition started at concentration of 10 g/l of urea. These results revealed that urea has both a positive and negative effect on growth of strain MB284.

There showing that despite the initial inhibitory effect of urea, the higher values for final biomass concentrations of solution happened in higher concentrations of urea. Therefore, in order to further investigate the toxic effects of urea on MB284, we examined the initial rate of urea degradation at different initial urea concentrations, under both aerobic and anoxic conditions (Figure 6). As shown in Figure 6, the rate of urea hydrolysis follows the Haldane effect, where at lower concentration rates of degradation increase followed by a rapid decline in rates after a certain concentration (approximately 3 g/L for aerobic conditions and 7 g/L for anoxic conditions). Also, Bhat et al reported a similar trend for urea hydrolysis by mixed consortia from loamy soil that follows Haldane kinetics [37].

**Figure 6:**
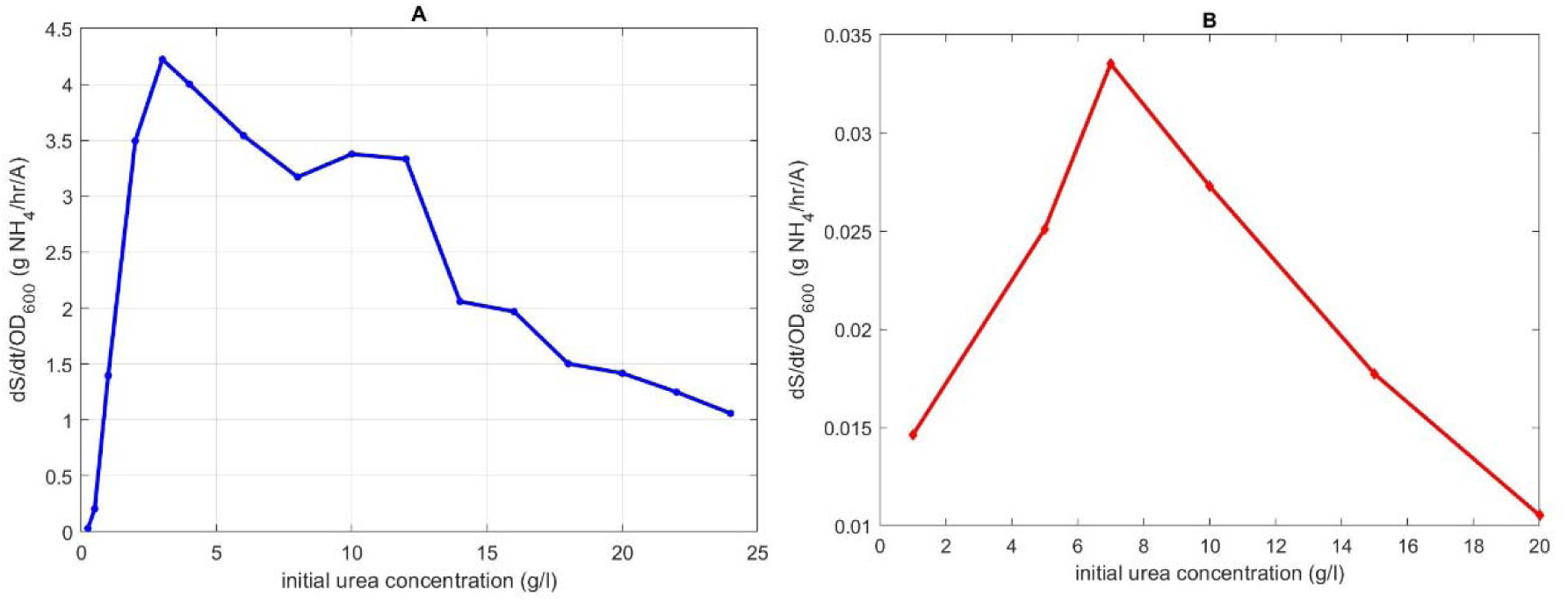
Halden equation in A: aerobic and B: anoxic conditions. Different initial concentration of urea were added to the MSM medium containing 10 g/l casamino acids.

Overall, the results of this study show that the components of the culture media, specifically the nutrient source and amount of urea, can have significant impacts on the growth rate and the MICP activity of *L. sphaericus* ATCC 13805TM (strain MB284). These results provide critical information on the growth requirements and growth behavior of strain MB284, especially in regard to use as a bioagent in the bio-consolidation of concrete.

## 4. Acknowledgment

This material is based upon work supported by the National Science Foundation (NSF) under Grant No 2029555, so we would like to appreciate NSF for its support.

## References

[1] A. I. Omoregie, E. A. Palombo and P. M. Nissom, “Bioprecipitation of calcium carbonate mediated by ureolysis: A review,” Environmental Engineering Research, vol. 26, no. 6, 2020.

[2] X. Zhu, J. Wang, N. D. Belie and N. Boon, “Complementing urea hydrolysis and nitrate reduction for improved microbially induced calcium carbonate precipitation,” Applied Microbiology and Biotechnology, 2019.

[3] A. C. Mitchell, E. J. Espinosa-Ortiz, S. L. Parks, A. J. Phillips, A. B. Cunningham and a. R. Gerlach, “Kinetics of calcite precipitation by ureolytic bacteria under aerobic and anaerobic conditions,” Biogeosciences, vol. 16, p. 2147–2161, 2019.

[4] P. Anbu, C. Kang, Y. Shin and J. So, “Formations of calcium carbonate minerals by bacteria and its multiple applications,” SpringerPlus, pp. 5–250, 2016.

[5] K. Chetty, S. Xie, Y. Song, T. McCarthy, U. Garbe, X. Li and G. Jiang, “Self-healing bioconcrete based on non-axenic granules: A potential solution for concrete wastewater infrastructure,” Journal of Water Process Engineering, vol. 42, 2021.

[6] Y. Su, T. Zheng and C. Qian, “Application potential of Bacillus megaterium encapsulated by low alkaline sulphoaluminate cement in self-healing concrete,” Construction and Building Materials, 2020.

[7] V. Achal and C. S. Chin, Building Materials for Sustainable and Ecological Environment, Springer Nature, 2021.

[8] L. Baumgartner, R. Reid, C. Dupraz, A. Decho, D. Buckley, J. Spear, K. Przekop and P. Visscher, “Sulfate reducing bacteria in microbial mats: Changing paradigms, new discoveries,” Sedimentary Geology, vol. 185, p. 131–145, 2006.

[9] M. J. Castro-Alonso, L.E. Montañez-Hernandez, M.A. Sanchez-Muñoz, M. R. M. Franco, R. Narayanasamy and N. Balagurusamy, “Microbially Induced Calcium Carbonate Precipitation (MICP) and Its Potential in Bioconcrete: Microbiological and Molecular Concepts,” Frontiers in Materrials, vol. 6, no. 126, 2019.

[10] H. Kim, H. Son, J. Seo and H. Lee, “Recent advances in microbial viability and self-healing performance in bacterial-based cementitious materials: A review,” Construction and Building Materials, vol. 274, 2021.

[11] W. D. Muynck, K. Verbeken, N. D. Belie and W. Verstraete, “Influence of temperature on the effectiveness of a biogenic carbonate surface treatment for limestone conservation,” Appl Microbiol Biotechnol, vol. 97, p. 1335–1347, 2013.

[12] C. Farrugia, R. P. Borg, L. Ferrara and J. Buhagiar, “The Application of Lysinibacillus sphaericus for Surface Treatment and Crack Healing in Mortar,” Front. Built Environ., vol. 5, no. 62, 2019.

[13] Y. S. Lee, H. J. Kim and W. Park, “Non-ureolytic calcium carbonate precipitation by Lysinibacillus sp. YS11 isolated from the rhizosphere of Miscanthus sacchariflorus,” Journal of Microbiolog, vol. 55, no. 6, p. 440–447, 2017.

[14] J. Intarasoontron, W. Pungrasmi, P. Nuaklongd, P. Jongvivatsakul and S. Likitlersuang, “Comparing performances of MICP bacterial vegetative cell and microencapsulated bacterial spore methods on concrete crack healing,” Construction and Building Materials, vol. 302, 2021.

[15] J. Wang, H. M. Jonkers, N. Boon and N. D. Belie, “Bacillus sphaericus LMG 22257 is physiologically suitable on self-healing concrete,” Applied Microbial and Physiology, 2017.

[16] F. G. Ferris, V. Phoenix, Y. Fujita and R. W. Smith, “Kinetics of calcite precipitation induced by ureolytic bacteria at 10 to 20°C in artificial groundwater,” Geochimica et Cosmochimica Acta, vol. 67, no. 8, p. 1701–1722, 2003.

[17] Y. Fujita, F. Ferri, R. Lawson, F. Colwell and R. Smith, “Subscribed Content Calcium Carbonate Precipitation by Ureolytic Subsurface Bacteria,” Geomicrobiology Journal, vol. 17, no. 4, pp. 305-318, 2000.

[18] D. J. Tobler, M. O. Cuthbert, R. B. Greswell, M. S. Riley, J. C. Renshaw, S. Handley-Sidhu and V. R. Phoenix, “Comparison of rates of ureolysis between Sporosarcina pasteurii and an indigenous groundwater community under conditions required to precipitate large volumes of calcite,” Geochimica et Cosmochimica Acta, vol. 75, p. 3290–3301, 2011.

[19] S. Stocks-Fischer, J. K. Galinat and S. S. Bang, “Microbiological precipitation of CaCO3,” Soil Biology and Biochemistry, vol. 31, pp. 1563–1571, 1999.

[20] H. Chen, Y. Huang, ·. C. Chen, J. P. Maity and C. Chen, “Microbial Induced Calcium Carbonate Precipitation (MICP) Using Pig Urine as an Alternative to Industrial Urea,” Waste and Biomass Valorization, vol. 10, p. 2887–2895, 2019.

[21] N. S. Adzami, M. F. Ghazali, A. H. Ramli, H. A. Tajarudin and Z. Daud, “A new potential of calcium carbonate production induced by Bacillus sphaericus in batch fermentation,” International Journal of Integrated Engineering, vol. 10, no. 9, pp. 130-134, 2018.

[22] M. S. b. Azmi, T. S. Lin, H. A. b. Tajarudin, M. b. M. Z. Makhtar and Z. b. Daud, “Characterisation of Calcium Carbonate Formed by Bacillus Sphaericus Via Fermentation of Urea,” International Journal of Integrated Engineering:, vol. 10, no. 9, pp 119–124, 2018.

[23] Y. C. Ersan, J. Y. Wang, N. Boon and N. D. Belie, “Ureolysis and denitrification based microbial strategies for self-healing concrete,” in Concrete Solutions, 2014.

[24] P. J. White and H. K. Lotay, “Minimal Nutritional Requirements of Bacillus sphaericus NCTC 9602 and 26 other Strains of this Species: the Majority Grow and Sporulate with Acetate as Sole Major Source of Carbon,” Journal of Geiierul Microbiolog, vol. 118, pp. 13–19, 1980.

[25] S. A. Rahmaninezhad, N. Mehrdadi and Z. Mahzari, “Analysis of the factors controlling the performance of a photoelectrocatalytic cell separated by UF membrane in degrading methylene blue,” Journal of the Australian Ceramic Society, vol. 57, no. 1, pp. 163–172, 2021.

[26] S. A. Rahmaninezhad, N. Mehrdadi and Z. Mahzari, “Comparison of the ultra-filtration and cation exchange membrane performance in photo electro catalytic degradation of methylene blue,” International Journal of Energy & Environment, vol. 10, no. 5, pp. 271–280, 2019.

[27] T. Zhu and M. Dittrich, “Carbonate Precipitation through Microbial Activities in Natural Environment, and Their Potential in Biotechnology: A Review,” Frontiers in bioengineering and biotechnology, vol. 4, 2016.

[28] P. Anbu, C. Kang, Y. Shin and J. So, “Formations of calcium carbonate minerals by bacteria and its multiple acteria and its multiple,” SpringerPlus, pp. 5–250, 2016.

[29] S.-F. S G. Jk and B. Ss, “Microbiological precipitation of CaCO3,” Soil Biol Biochem, vol. 31, p. 1563–157, 1999.

[30] I. S. Tan and K. S. Ramamurthi, “Spore formation in Bacillus subtilis,” Environ Microbiol Rep., p. 212–225, 2014.

[31] C. Akali and A. YetıŞemıyen, “Use of whey powder and skim milk powder for the production of fermented cream,” Food Science and Technology, vol. 36, no. 4, pp. 616–621, 2016.

[32] L. A. v. Paassena, C. M. Daza, M. Staal, D. Y. Sorokin, W. v. d. Zon and M. C. v. Loosdrecht, “Potential soil reinforcement by biological denitrification,” Ecological Engineering, vol. 36, p. 168–175, 2010.

[33] A. Yoneyama, H. Choi, M. Inoue, J. Kim, M. Lim and Y. Sudoh, “Effect of nitrite/nitrate-based accelator on the strength development and hydrate formation in cold-weather cementitious materials,” Materials, vol. 14, no. 1006, 2021.

[34] B. C. J. G. Knight and H. Proom, “A Comparative Survey of the Nutrition and Physiology of Mesophilic Species in the Genus Bacillus,” Journal of General Microbiology, vol. 4, pp. 508–538, 1950.

[35] G. H. Bornside and R. E. Kallio, “Urea-hydrolyzing Bacilli II. Nutritional profiles,” Journal of Bacteriology, vol. 71, pp. 655–660, 1956.

[36] M. Neumann-Schaal, J. D. Hofmann, S. E. Will and D. Schomburg, “Time-resolved amino acid uptake of Clostridium difficile 630Δerm and concomitant fermentation product and toxin formation,” BMC Microbiology, vol. 15, no. 281, 2015.

[37] M. R. Bhat, D. V. R. Murthy and M. B. Saidutta, “Urea Hydrolysis in Saturated Loam Soil,” Asian Research Publishing Network (ARPN), vol. 6, no. 3, 2011.

